# Rapid divergence of the copulation proteins in the *Drosophila dunni* group is associated with hybrid post-mating-prezygotic incompatibilities

**DOI:** 10.1101/2020.05.20.106724

**Authors:** Tom Hill, Hazel-Lynn Rosales-Stephens, Robert L. Unckless

## Abstract

Proteins involved in post-copulatory interactions between males and females are among the fastest evolving genes in many species and this has been attributed to reproductive conflict. Likely as a result, these proteins are frequently involved in cases of post-mating-prezygotic isolation between species. The *Drosophila dunni* subgroup consists of a dozen recently diverged species found across the Caribbean islands with varying levels of hybrid incompatibility. We sought to examine how post-mating-prezygotic factors are involved in isolation among members of this species group. We performed experimental crosses between species in the *dunni* group and find evidence of hybrid inviability. We also find an insemination reaction-like response preventing egg laying and leading to reduced female survival post-mating. To identify that genes may be involved in these incompatibilities, we sequenced and assembled the genomes of four species in the *dunni* subgroup and looked for signals of rapid evolution between species. Despite low levels of divergence, we found evidence of rapid evolution and divergence of some reproductive proteins, specifically the seminal fluid proteins. This suggests post-mating-prezygotic isolation as a barrier for gene flow between even the most closely related species in this group and seminal fluid proteins as a possible culprit.

## Introduction

Numerous groups of recently diverged species have been used to study speciation across multicellular taxa (Coyne and Orr 1989; Mckinnon and Rundle 2002; Glor *et al*. 2005; Kitano *et al*. 2009; Brekke and Good 2014). These and other studies find an array of complex relationships between species caused by varying levels of divergence across genomes, incomplete isolation and differing forms of reinforcement (Coyne and Orr 2004; Orr 2004; Presgraves 2007; Matute *et al*. 2010; Moyle and Nakazato 2010; Orr *et al*. 2013; Payseur and Rieseberg 2016). Recently diverged species with incomplete reproductive barriers prove to be more useful for understanding how new species can evolve (Coyne and Orr 1989; GourbiÈre and Mallet 2010; Presgraves 2010). These species groups can be used in QTL studies to identify loci which contribute to the reduced fitness of hybrids (Howard *et al*. 2002; Noor *et al*. 2007; Kitano *et al*. 2009), or to identify genes which may be involved in the early stages of speciation, such as those causing inviability or sterility in the heterogametic sex (a phenomenon known as Haldane’s Rule) (Haldane 1922; Coyne and Orr 1989; Orr 1995; Gavrilets and Waxman 2002; Coyne and Orr 2004; Orr 2013).

Several studies have also highlighted that proteins transmitted in the seminal fluid to the female reproductive tract may also drive isolation as a post-mating-prezygotic mating barrier for incompletely separated species, either caused by, or resulting in, reinforcement (Gavrilets and Waxman 2002; Coyne and Orr 2004; Andres *et al*. 2008; GourbiÈre and Mallet 2010; Larson *et al*. 2012; Larson *et al*. 2013; AHMED-BRAIMAH 2016; Turissini *et al*. 2017; Matute *et al*. 2020). Barriers to hybridization have also been examined in different *Drosophila* species groups, finding varying levels of divergence, and in some cases the mechanisms for isolation between species (Patterson 1947; Grant 1983; Coyne and Orr 1989; Presgraves 2007; Miller *et al*. 2010; Matute and Ayroles 2014; AHMED-BRAIMAH 2016; Turissini *et al*. 2017; Matute *et al*. 2020). Some studies, focusing on the effects of heterospecific matings on females, have found drastic changes in the females, including the swelling of the reproductive tract (Patterson 1947) and the activation of stress response pathways (AHMED-BRAIMAH *et al*. 2020), likely due to antagonistic interactions between male seminal fluid proteins that the heterospecific female tract (Knowles and Markow 2001). These responses likely result in reinforcement of diverging reproductive behavior to prevent such matings (Coyne and Orr 2004; Turissini *et al*. 2017).

The *Drosophila dunni* subgroup is found within the *cardini* group in the *Drosophila* subgenus (Supplementary Figure 1) (Heed 1962). This species group diverged across the Caribbean islands thousands of years ago creating endemic populations, each on a different island or set of islands (Heed 1962; Hollocher *et al*. 2000; Wilder and Hollocher 2003). Despite their extended isolation from each other, species are still able to hybridize (to varying levels of success) and are a useful species group for understanding several traits, such as the evolution of pigmentation or reproductive isolation (Stalker and Streisinger 1953; Patterson 1954; Hollocher *et al*. 2000; Wilder and Hollocher 2003). In some cases, these hybrid offspring show evidence of Haldane’s rule (Haldane 1922; Orr 2013), with crosses producing only female offspring, or sterile male offspring (Heed 1962).

**Figure 1:**
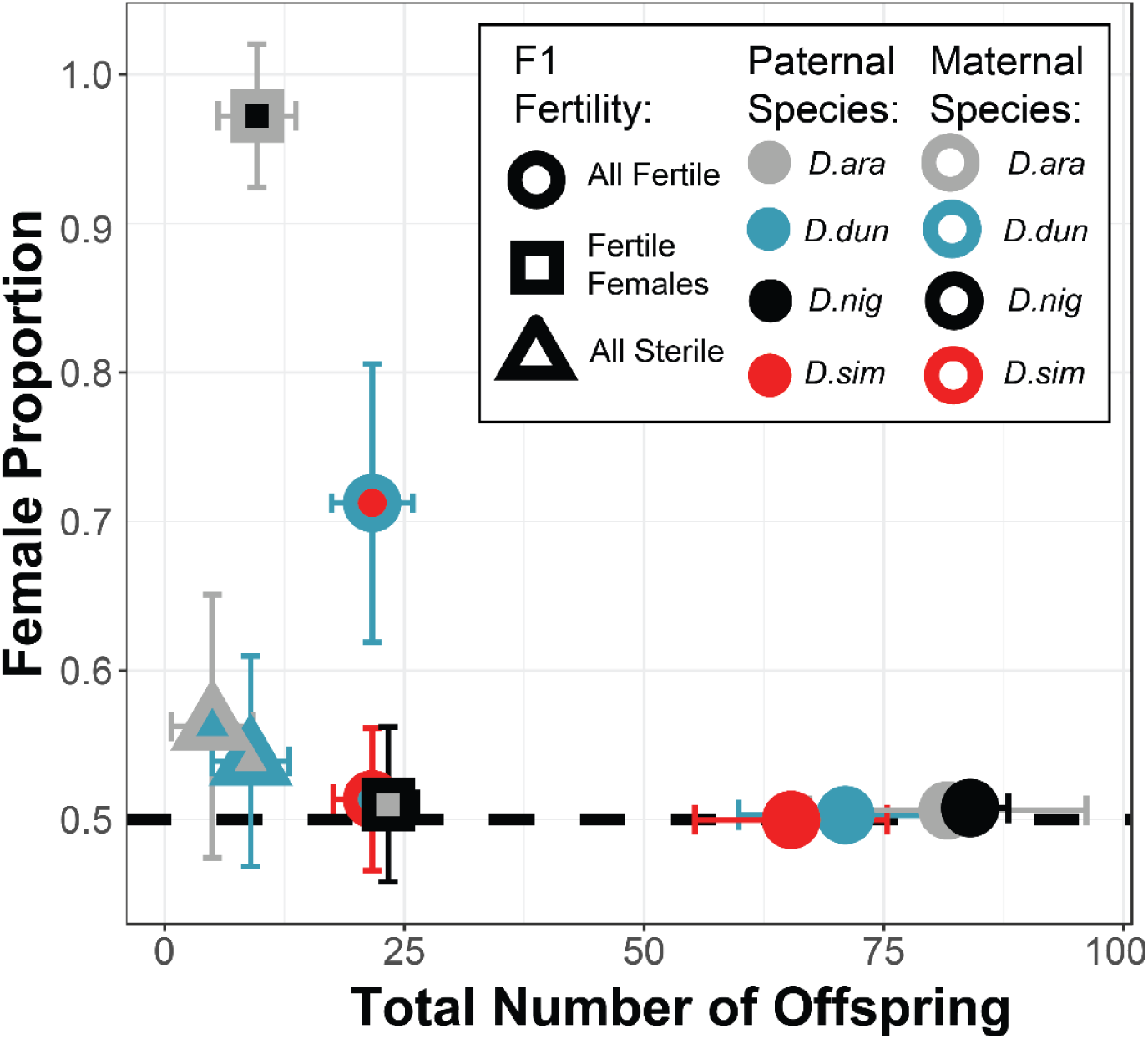
Mean number of offspring produced by three replicates of 10 females of each species when crossed to males of different species. Points of the same color represent conspecific crosses while dots with a different center represent a cross between two different species. Point shape shows the state of fertility of F1 offspring, either both fertile, both sterile or only females fertile. Error bars show the standard deviation of offspring count and sex ratio across replicates. *D. ara* = *D. arawakana, D. dun* = *D. dunni, D. nig* = *D. nigrodunni, D. sim* = *D. similis*. While all we performed all pairwise heterospecific crosses, only crosses which produced offspring are shown on the plot.

Here we perform experimental crosses in the *dunni* group and find that in some crosses, heterospecific matings reduces female survival compared to conspecific matings, potentially caused by an insemination reaction-like effect (Patterson 1947). Using a combination of long-read and short-read sequencing, we assembled the genomes of four species in the *dunni* group to identify proteins driving this incompatibility. We find these genomes are of similar quality and composition as other higher quality genomes in the *Drosophila* subgenus (Zhou *et al*. 2012; Zhou and Bachtrog 2015; Gramates *et al*. 2017; Hill *et al*. 2019). We also estimate rates of evolution across these genomes and identify several pathways of groups of genes of interest diverging between species (particularly between *D. nigrodunni* and *D. arawakana*), such as a divergence in immune pathways and in seminal fluid proteins.

## Materials and Methods

### Drosophila stocks, experimental crosses and survival assays

We obtained stocks for *Drosophila arawakana* (stock number: 15182-2260.00), *D. dunni* (stock number: 15182-2291.00), *D. nigrodunni* (stock number: 15182-2311.00) and *D. similis* (stock number: 15182-2321.00) from the Cornell *Drosophila* species stock center. Each species was maintained on standard instant fly food (Formula 4-24, Carolina Biological Supply Company, Burlington, NC) in an incubator at 23°C. Before experiments, we inbred for three generations. Specifically, we established 10 single fly crosses for each species and chose a single successful cross per generation. We then repeated this for three generations. We then randomly chose one inbred vial to work with for the remainder of the experiments described.

### Experimental crosses within and between species

We performed initial crosses in all pairwise combinations of species, for both directions of the cross, as well as within species crosses, to confirm previous assessments of between species viability (Heed 1962; Wilder and Hollocher 2003).

For each species we cleared vials of adults at 9:00AM central time and collected any emerged adults in 3-hour intervals following this, separating by sex. We then used these virgin flies to mate all species in pairwise combinations in 3 replicates. For each replicate we mated 10 males with 10 females (all aged 2-3 days) for 5 days (Wilder and Hollocher 2003; Cenzi de RÉ *et al*. 2010). We then collecting offspring every day for 30 days following the removal of the parents. After aging virgin F1 offspring for 3 days, we separated these into groups of 10 flies of the same parental species and mated with 10 flies of the opposite sex (5 the paternal species, 5 the maternal species) to assess the fertility of the F1 flies. As *D. arawakana* appeared to be infected with *Wolbachia*, we sought to cure all species of any bacteria which may affect crosses. We created sublines of each species raised on food containing tetracycline-hydrochloride (0.05mg/ml) for three generations. Following this, we extracted DNA from females of each strain and tested for *Wolbachia* using PCR (wsp-81F (5′-TGGTCCAATAAGTGATGAAGAAAC-3′), wsp-691R (5′-AAAAATTAAACGCTACTCCA-3′), producing a ∼600bp product from 10uL reactions, under the following cycling conditions: 94°C for 4 min, followed by 30 cycles of 40 s at 94°C, 40 s at 55°C, 1 min at 72°C and a final extension step of 10 min at 72°C) (Zhou *et al*. 1998). We then repeated experimental crosses, as described above, with the tetracycline cured strains.

We assayed female survival for *D. arawakana, D. dunni, D. nigrodunni* and *D. similis* in virgins and following mating, in both uncured and tetracycline cured flies. We considered a cross to be conspecific if we mated within species and a cross to be heterospecific if we mated with the most closely related species where fertile hybrids were found in previous crosses (e.g. *D. dunni* to *D. similis* and *D. arawakana* to *D. nigrodunni*). For these crosses we established 5-15 vials of 10 males and 10 females of the given species (with no males when measuring virgin females), all aged 2-3 days. We then recorded the survival of females every day (checking at 10AM Central time) for 30 days, flipping the flies onto new food every 3-4 days and removing males after the first 5 days. We then fit a survival curve across the total data for each cross type using SurvMiner (Kassambara *et al*. 2017) in R (R-CORE-TEAM 2013) and used a Cox’s Hazard Ratio to identify significant differences in survival between sets of crosses. For the initial crosses we used the following model:

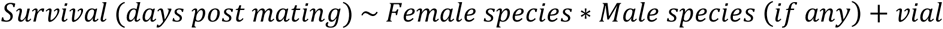

We set the reference level as the conspecific cross (e.g. *D. arawakana ♂ × D. arawakana* ♀) and looked for significant differences from these for interaction terms to determine if unmated females (e.g. *D. arawakana* ♀ not mated) or heterospecifically crossed females (e.g. *D. arawakana ♂ × D. nigrodunni* ♀) show significant differences from the conspecific cross. To consider the effect of *Wolbachia* infection on these crosses, we repeated these initial crosses alongside the same crosses with *Wolbachia* cured flies (cured as described above) and a Cox’s Hazard Ratio was used to determine the effect of *Wolbachia* on survival, and to test for differences in survival between sets of crosses after accounting for *Wolbachia*:

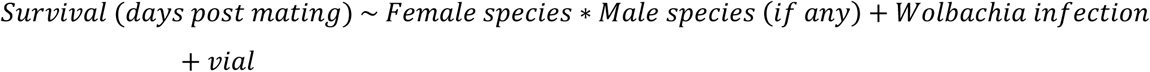

### Post-mating dissection of the female reproductive tract

We collected virgin males and females for tetracycline-cured *D. arawakana* and *D. nigrodunni* as described above and aged them 2-3 days. We then established conspecific and heterospecific experimental crosses for 6 replicates of 10 males and 10 females at 10AM central time, as well as virgin control females for 6 replicates of 10 females. Following 24 hours of cohabitation, for 3 replicates of each cross, we separated the females for each cross and dissected the reproductive tract. Based on previous work describing the insemination reaction (Patterson 1947; Grant 1983; Markow and Ankney 1988), we scored the reproductive tract for each female, identifying if the female had mated (by the presence of sperm), if the reproductive tract appeared to be swollen (relative to the unmated virgin females) or if the reproductive tract was destroyed or damaged (alongside a swollen tract, if possible to tell). We repeated this scoring for the remaining 3 replicates of each cross 24 hours later (48 hours total). We then compared conspecific and heterospecific crosses for rates of mating and rates of insemination reaction occurrence.

### Genome sequencing, assembly and annotation

We extracted DNA following the protocol described in (Chakraborty et al. 2017) for *D. arawakana, D. dunni, D. nigrodunni* and *D. similis* females. We prepared the *D. dunni* and *D. nigrodunni* DNA as a sequencing library using the Oxford Nanopore Technologies Rapid 48-hour (SQK-RAD002) protocol, which we then sequenced separately using a MinION (Oxford Nanopore Technologies, Oxford, UK) (Jain *et al*. 2016) (Supplementary Table 1). We also prepared the *D. arawakana, D. dunni, D. nigrodunni* and *D. similis* samples as Illumina libraries with a 300bp insert size which we sequenced on an Illumina HiSeq4000 to produce 150bp paired-end reads (Supplementary Table 1). We removed Illumina adapters using Sickle (Joshi and Fass 2011) and trimmed the Illumina sequences using Scythe (Buffalo 2018). For the two MinION genomes, bases were called *post hoc* using the built in read_fast5_basecaller.exe program with options: –f FLO-MIN106 –k SQK-RAD002 –r–t 4. For *D. dunni*, raw reads were assembled using Minimap2 and Miniasm (parameters: -x ava -o nt -t 8) (Li 2016). We then polished using Racon with Oxford Nanopore Technology reads for three iterations and Pilon with Illumina fragment library reads for three iterations (Walker *et al*. 2014). For the *D. nigrodunni* genome, we first used wtdbg2 to assemble the genome (parameters: -t 4 -L 1000) (Ruan and Li 2020). We then created a second assembly using Minimap2. For each, we ran Racon and Pilon for three iterations as described for *D. dunni*, then merged the two *D. nigrodunni* assemblies using Quickmerge (Liu and Yang 2013). Following this, we polished this merged genome using Pilon for four more iterations. Both assemblies were benchmarked using BUSCO (v 3.0.2) and the *Diptera* database (SIMÃO *et al*. 2015).

For *D. similis*, we mapped data to the *D. dunni* genome before Pilon polishing and polished the *D. dunni* genome using *D. similis* data in Pilon for three iterations, to insert *D. similis* variants into the genome. Following this we mapped *D. similis* data to this genome using BWA (Li and Durbin 2009) and SAMtools (LI *et al*. 2009), and called variants using Picard (BROAD-INSTITUTE 2017) and GATK Haplotypecaller (Mckenna *et al*. 2010; Depristo *et al*. 2011). We then used BCFtools (Narasimhan *et al*. 2016) to filter these variants, removing calls below a quality threshold of 200 and inserted them into the polished genome. This was repeated for two more iterations to create a *D. similis* alternate genome. The same pipeline was followed for *D. arawakana* mapped to the *D. nigrodunni* genome.

We used the *D. innubila* transcriptome (Hill *et al*. 2019) as well as protein databases from *D. innubila, D. virilis, D. melanogaster*, and *M. domestica* in MAKER2 (Holt and Yandell 2011) to annotate each genome, including using *RepeatModeler* (Smit and Hubley 2008) in an attempt to correctly assign repetitive regions and retraining a HMM using SNAP following each iteration (Johnson *et al*. 2008). This was repeated for three iterations to generate a GFF file containing gene evidence generated by MAKER2 (Holt and Yandell 2011).

Finally, we identified orthologous genomic regions pairwise for each of the four species examined here to each other and to the *D. innubila* genome using progressiveMauve (Darling *et al*. 2004). We visualized orthologous regions using rCircos (Zhang *et al*. 2013). We attempted to confirm any apparent structural differences based on progressiveMauve by mapping short reads for each species to a different genome and calling copy number differences using Delly (Rausch *et al*. 2012) and dudeML (Hill and Unckless 2019), taking the consensus of the two tools, but favoring the absence of a copy number variant when we found discrepancies between the two tools.

### Assessing the repetitive content across the dunni group

For each genome, we identified the repetitive content *de novo* using RepeatModeler to call the repeats (engine = NCBI) (Smit and Hubley 2008) and RepeatMasker (-gff –gcalc –s) to identify the repetitive regions (Smit and HUBLEY 2013-2015). We also used dnaPipeTE (genome coverage = 1, sample number = 2, cpu = 4, genome size = 168000000) (Goubert *et al*. 2015) to identify the repetitive content in the short-read data for each species, which we used to make a second map of reference genome repetitive regions using RepeatMasker. For both sets of repeat content assemblies we identified which TE families were shared between species and which were unique to species using blastn (e-value < 10e-5, hsps = 1, alignments = 1). We then identified what proportion of the genome each TE family constituted across species.

### Placing the dunni group in the Drosophila phylogeny

To find the consensus species tree despite the differing evolutionary histories of different genes (Mendes and Hahn 2016), we randomly sampled 100 genes conserved across *Drosophila* and humans from and extracted these from our four focal species, as well as from several other *Drosophila* species, taken from Flybase (Gramates *et al*. 2017) and the NCBI genomes database (Zhou *et al*. 2012; Hamilton *et al*. 2014; Palmieri *et al*. 2014; Zhou and Bachtrog 2015; Kitts *et al*. 2016; Hill *et al*. 2019). We then aligned each gene group separately using MAFFT (--auto) (Katoh *et al*. 2002) and created a multiple gene super-tree based on the consensus of each gene tree, following 100 bootstraps with PhyML (-b 100 -N 100 -GTR -gamma 8) (Le and Gascuel 2008; Guindon *et al*. 2010). We also generated gene trees for each of the 100 genes independently, following the same protocol. In this case 66 of the 100 trees gave the same topology of the *dunni* group as the total tree, while 7 trees had distinct topologies and 27 trees gave the topology of *D*.*similis* as an outgroup to the other three species, with *D. dunni* a sister to the *D. nigrodunni-D*.*arawakana* complex.

### Estimating rates of evolution across the dunni group

For each gene in the genomes of our four focal species, we identified orthology to each other and to genes in *D. innubila* using blastp (e-value < 0.00001, hsp = 1 alignment = 1) (Altschul *et al*. 1990). For each set of orthologs, we aligned using PRANK to generate a codon alignment and gene-tree (LÖYTYNOJA 2014), as subtle differences between the species tree and gene trees can result in false estimates of divergence (Mendes and Hahn 2016). We then estimated rates of both non-synonymous and synonymous substitutions using codeML (Yang 2007), we estimated specific rates of evolution along each branch of the *dunni* group and leading into the *dunni* group using *D. innubila* as an outgroup (model 0) (Yang 2007). Specifically, we estimated synonymous divergence (dS), non-synonymous divergence (dN) and the proportion of the two values (dN/dS). Finally, we also estimated rates of evolution across the entire *dunni* group phylogeny using codeML (models 7 & 8) (Yang 2007), choosing the best fitting model using a likelihood ratio test (*p*-value < 0.05).

Using the estimated rates of evolution, we then compared the rates of evolution across the entire phylogeny and on specific branches to each species, for genes of similar levels of synonymous divergence (dS, windows of 0.001 dS, e.g. all genes within 0.001 dS of each other) we found the 97.5^th^ upper percentile for dN/dS. For the closely related species pairs (*D. nigrodunni* and *D. arawakana, D. dunni* and *D. similis*) we compared measures of dN/dS between species and found the 97.5^th^ upper percentile for dN/dS per species per window of dN/dS for the paired species (0.001, sliding 0.001).

We then took outlier genes (e.g. genes above the 97.5^th^ percentile in each category) and looked for enrichments in gene ontology categories compared to non-outlier genes using GOrilla (Eden *et al*. 2009). For GO categories of interest, such as those enriched for duplications or for high levels of dN/dS, we compared dN/dS of genes in these categories to the nearby genomic background. For each gene we extracted nearby genes (within 100kbp up or downstream on the same chromosome), of similar divergence levels on each branch (within 0.01 dS), we then found the difference in dN/dS between the median of the background genes and the focal gene. We then used a Wilcoxon-Rank Sum test to identify GO categories on each branch with significantly higher (or lower) dN/dS than the background.

Using the annotations of all species and *D. innubila*, we identified genes with more than one copy in one species, relative to all other species. We confirmed this by estimating copy numbers of genes in each species using short read information and dudeML (following the tutorial pipeline for N = 1) with the short read information mapped to the genome of the sister species (Hill and Unckless 2019). We then used GOrilla (EDEN *et al*. 2009) to identify Gene ontology categories that are enriched for duplicates on specific branches, which we confirmed using Panther (Thomas *et al*. 2003).

### Statistics

We used R for all statistics in this analysis (R-CORE-TEAM 2013), and ggplot2 for data visualization and figure production (Wickham 2009).

## Results

### The Drosophila dunni group shows varying levels of hybrid compatibility

The *Drosophila dunni* group is a species group endemic to islands in the Caribbean, with each island inhabited by a different complement of species (Stalker and Streisinger 1953; Wilder and Hollocher 2003; Cenzi de RÉ *et al*. 2010). These species have varying levels of hybrid incompatibilities, with some crosses producing viable offspring (e.g. *D. dunni* × *D. similis*) and others producing sterile offspring (e.g. *D. arawakhana* × *D. dunni*) or no offspring (e.g. *D. nigrodunni* × *D. similis*). In keeping with Haldane’s rule (Haldane 1922), some produce sterile males, or no males at all (Figure 1, Supplementary Table 2, e.g. *D. nigrodunni* × *D. arawakhana*). Despite divergence on levels comparable to the *D. melanogaster* subgroup (Supplementary Figure 1, Supplementary Table 3), there are no characterized inversions between species (Stalker and Streisinger 1953; Cordeiro *et al*. 2014), allowing differences across the species group to be investigated with a higher resolution than the *D. melanogaster* group allows.

Given the variety in levels of divergence and isolation between species, we examined the differences in this species group and identify patterns of divergence between species that could be associated with the reproductive isolation. Our focus is on the two hybrid crosses which produce some compatible offspring, such as with *D. nigrodunni* and *D. arawakana*, in which one direction of the heterospecific cross produces only female offspring (Figure 1).

### Drosophila arawakana males reduce the lifespan of D. nigrodunni females

We next determined if there was evidence of further effects, beyond offspring viability (precopulatory, prezygotic and postzygotic) on hybridization. To do this, we established crosses between species, focusing on crosses that produced some fertile offspring (*D. nigrodunni ♀ × D. arawakana ♂, D. arawakana ♀ × D. nigrodunni ♂, D. similis ♀ × D. dunni ♂, D. dunni ♀ × D. similis ♂*, Supplementary Table 2). We also established matched crosses within species, and a matched control of virgin females. For each cross we recorded the survival of females following 5 days of mating.

In all cases, and consistent with studies in *D. melanogaster* (Chapman et al. 1993; Wigby and Chapman 2005), virgins generally survive longer than mated females, though not significantly in some cases (Figure 2, Cox Hazard Ratio z-value = 3.868, *p*-value = 0.00011). The heterospecific crosses showed no difference from the conspecific crosses for *D. similis* and *D. dunni* (Figure 2, Cox Hazard Ratio z-value = −0.488, *p*-value = 0.62545), though *D. similis* heterospecifically mated females lived longer than conspecifically mated females (Figure 2, Cox Hazard Ratio z-value = 2.153, *p*-value = 0.03134). In contrast, when *D. nigrodunni* females are crossed to *D. arawakana* males, females have significantly decreased survival compared to conspecific crosses and virgin females (Figure 2, Cox Hazard Ratio z-value = −3.360, *p*-value = 0.00078), the same cross which also produced only female offspring (Figure 1).

**Figure 2:**
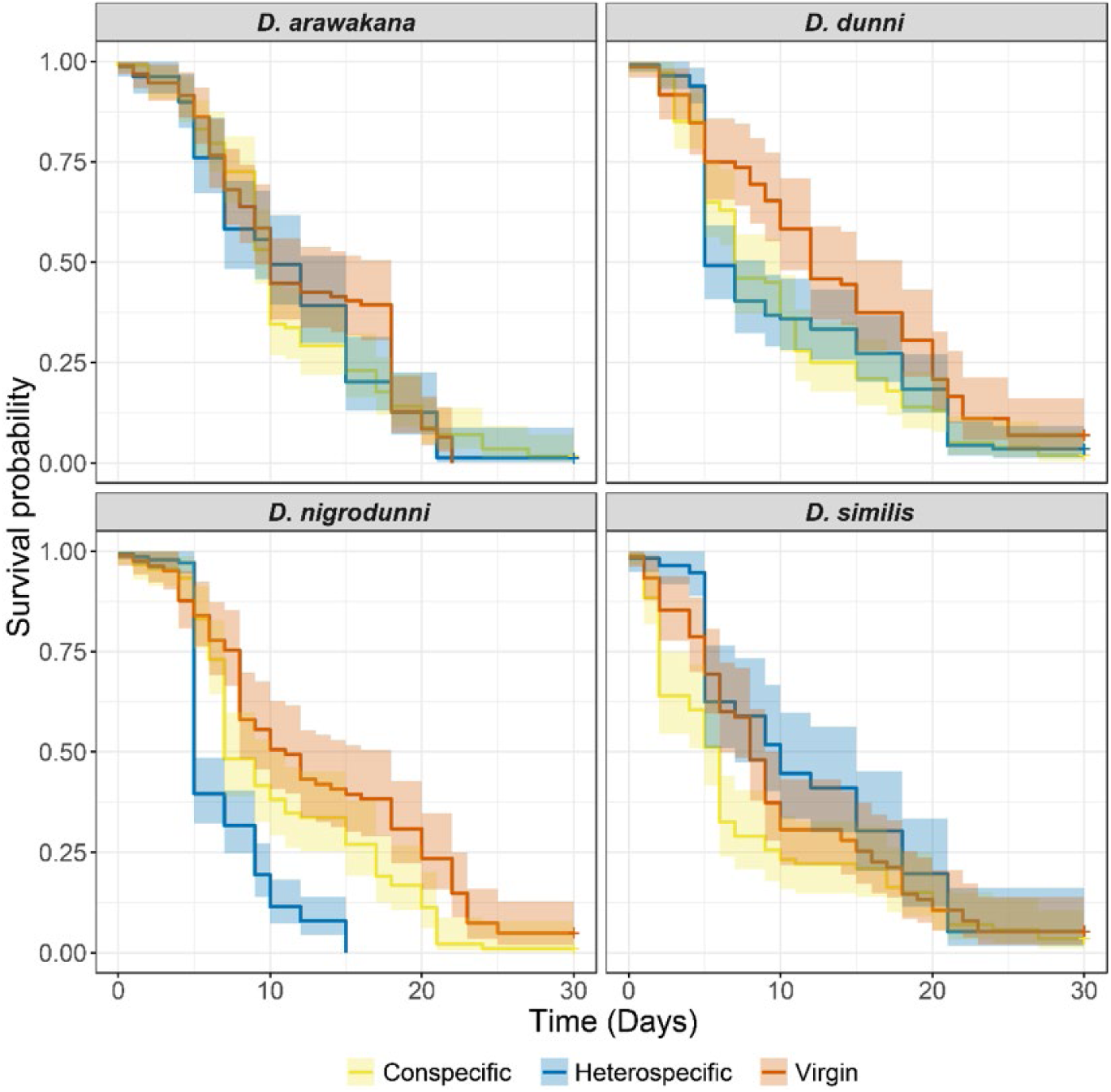
Survival of females postmating. Survivial probability of females for each species used in each cross, compared to virgin female survival. Females are separated by species, and grouped as virgins, conspecific crossed (crossed to own species), heterospecific crossed (crossed to a different species). In the case of heterospecific crosses, *D. arawakana* is only crossed to *D. nigrodunni* and *D. dunni* is only crossed to *D. similis*.

As the *D. arawakana* strain examined was infected with *Wolbachia* and the *D. nigrodunni* was not, we cured all strains of bacteria using tetracycline-hydrochloride and repeated the survival assays. All females in this second block have reduced survival compared to the original survival assay, (Supplementary Figure 2, Cox Hazard Ratio z-value = −5.654, *p*-value = 1.56e-08), suggesting a difference in the two experiments that could be attributed to Tetracycline-Hydrochloride exposure. In the tetracycline exposed flies, we again find reduced survival in the *D. nigrodunni* ♀ *x D. arawakana ♂* cross compared to the conspecific crosses (Supplementary Figure 3, Cox Hazard Ratio z-value = −3.815, *p*-value = 0.000136).

**Figure 3:**
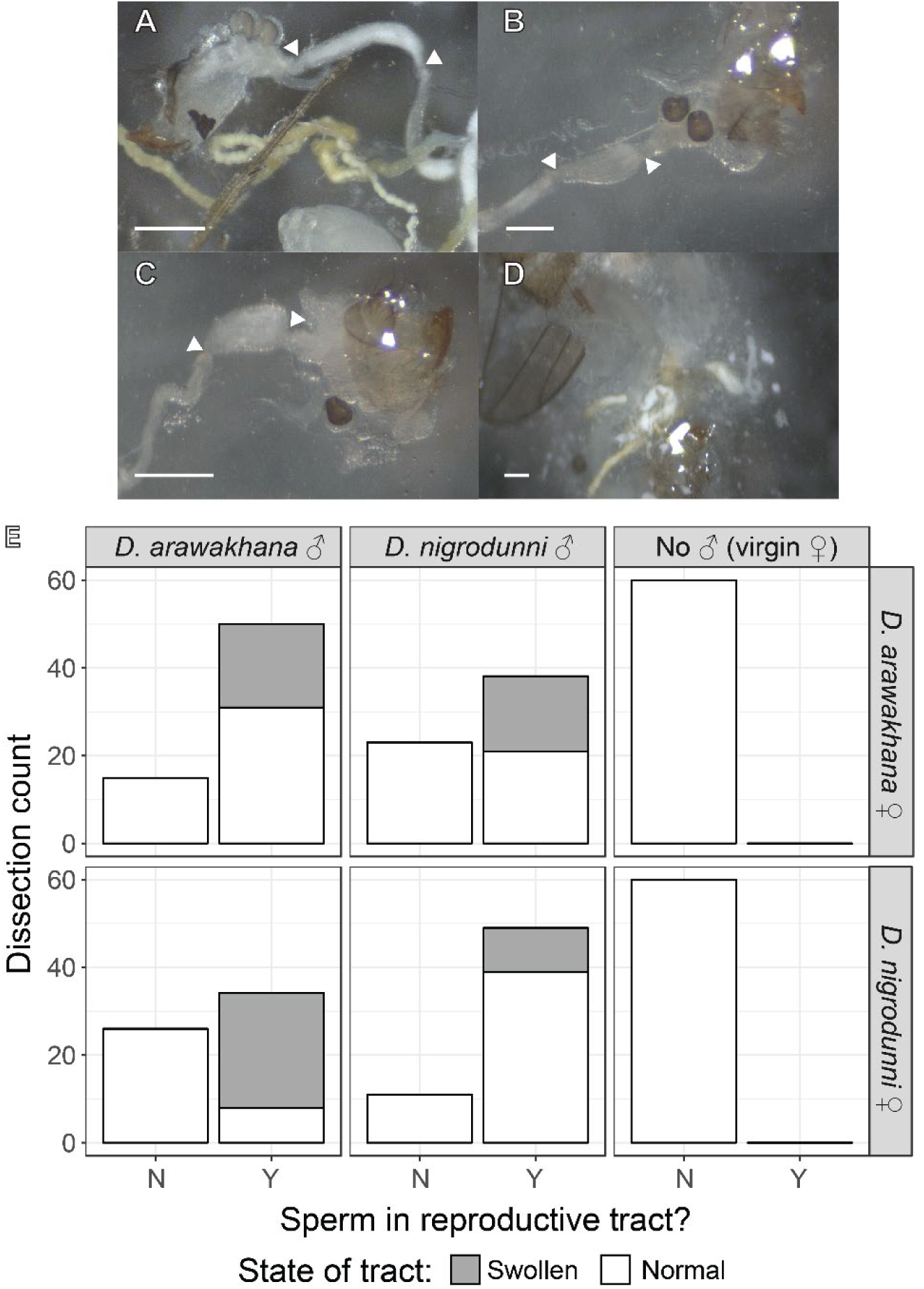
Abnormal insemination reactions may be responsible for reproductive isolation. **A-C**. Dissections showing differing conditions of the female reproductive tract. When applicable, arrows label the start and end of same section of the oviduct between dissections. Ovipositors and scale bar also shown for scale. **A**. Normal oviduct containing sperm. **B**. Normal oviduct with no sperm. **C**. Swollen oviduct containing sperm. **D**. Ruptured oviduct in sample with reaction mass-like phenotype. **E**. Plots summarizing rate of mating, and the effect of mating on the reproductive tract in crosses within and between *D. arawakana* and *D. nigrodunni*. Plots are separated by the male involved in the cross (columns) and the female involved in the cross (rows), with plots scoring the number of females with sperm in the reproductive tract, and if the tract was normal or swollen/damaged.

### The insemination reaction may be associated with the reduced female survival and reduced number of hybrid offspring

In several other hybrid crosses between species in the *Drosophila* subgenus of *Drosophila*, other studies have highlighted a reaction between the seminal fluid of one species with the environment of the reproductive tract in the other species, called the insemination reaction (Patterson 1947; Grant 1983; Markow and Ankney 1988). In the hours following mating, the reproductive tract swells, and, in some cases, proteins in the seminal fluid cause the formation of a “reaction mass”, a large dark mass which can burst through the wall of the tract (Patterson 1947).

Given the reduced survival of *D. nigrodunni* females following mating with *D. arawakana* males and the reduced number of hybrid offspring, we hypothesized that an incompatibility between the diverged seminal fluid proteins and the heterospecific reproductive tract could cause an abnormally deleterious reaction mass which reduces female survival.

We established experimental crosses within and between *D. arawakana* and *D. nigrodunni*. Then, 24 and 48 hours after crossing we dissected the females to identify whether sperm was present in the female reproductive tract (Figure 3A and B), and score for abnormal reproductive tracts consistent with the insemination reaction (Figure 3C and D). Interestingly, there was no significant differences between the number of mated females 24 and 48 hours after establishing crosses (Logistic regression: sperm presence ∼ collection date: z-value = 1.285, *p*-value = 0.198873), but did score significantly fewer mated females in heterospecific crosses versus conspecific crosses (Logistic regression: sperm presence ∼ cross type: z-value = −2.948, *p*-value = 0.00319). In several mated females when compared to virgin females, we find a swelling of the reproductive tract consistent with the insemination reaction (Figure 3C). Exclusively in several heterospecifically crossed females, we also saw damaged and destroyed reproductive tracts (Figure 3D). We find a significant excess of swollen/damaged tracts in heterospecifically mated *D. nigrodunni* compared to conspecific controls (Figure 3E, Logistic regression: swollen tract ∼ *D. nigrodunni* cross type: z-value = 4.723, *p*-value = 2.32e-06). While we do find swollen tracts in *D. arawakana* females we find no difference between heterospecific and conspecific females (Figure 3E, Logistic regression: swollen tract ∼ *D. arawakana* cross type: z-value = 0.493, *p*-value = 0.622162).

### Genes involved in copulation and immune defense have high rates of divergence between species

We reasoned that these incompatibilities between species could be caused by a divergence in copulation proteins. Previous work has suggested that females may be susceptible to bad reactions following hybrid matings due to no protection from the other species accessory gland proteins (Markow and Ankney 1988; Knowles and Markow 2001). Specifically, that there is an arms race between sexes to block/unblock the female reproductive tract and that females of other species have not evolved to suppress these reactions. Based on this, we sought to examine the levels of divergence and identify rapidly evolving genes between species. We sequenced, assembled and annotated the genomes of each species involved (see Materials and Methods), producing two high quality genomes with high synteny to each other and to *D. innubila* (Supplementary Tables 1 & 4 and Supplementary Figure 4A), and two assemblies derived from these *de novo* assemblies. The two *de novo* assemblies had high BUSCO scores (*D. dunni* scored 93.9%: 2627 complete, 79 fragmented and 93 missing out of 2799 total; *D. nigrodunni* scored 97.3%: 2721 complete, 37 fragmented and 41 missing out of 2799 total). Consistent with previous findings we find no large structural rearrangements between genomes, and no evidence of fixed inversions between species in the *dunni* group (Heed 1962; Cordeiro *et al*. 2014), though we do find several inversions between the next closest whole genome available, *D. innubila* on Muller elements B, C and D (*D. nigrodunni* shown in Supplementary Figure 4B). We annotated the *dunni* group genomes using a transcriptome from *D. innubila* in MAKER (Holt and Yandell 2011) and found between 10752 and 11581 genes in each species, most of which show orthology to previously identified genes in *D. virilis, D. melanogaster* or *D. innubila* (Supplementary Table 5) (Hill *et al*. 2019).

**Figure 4:**
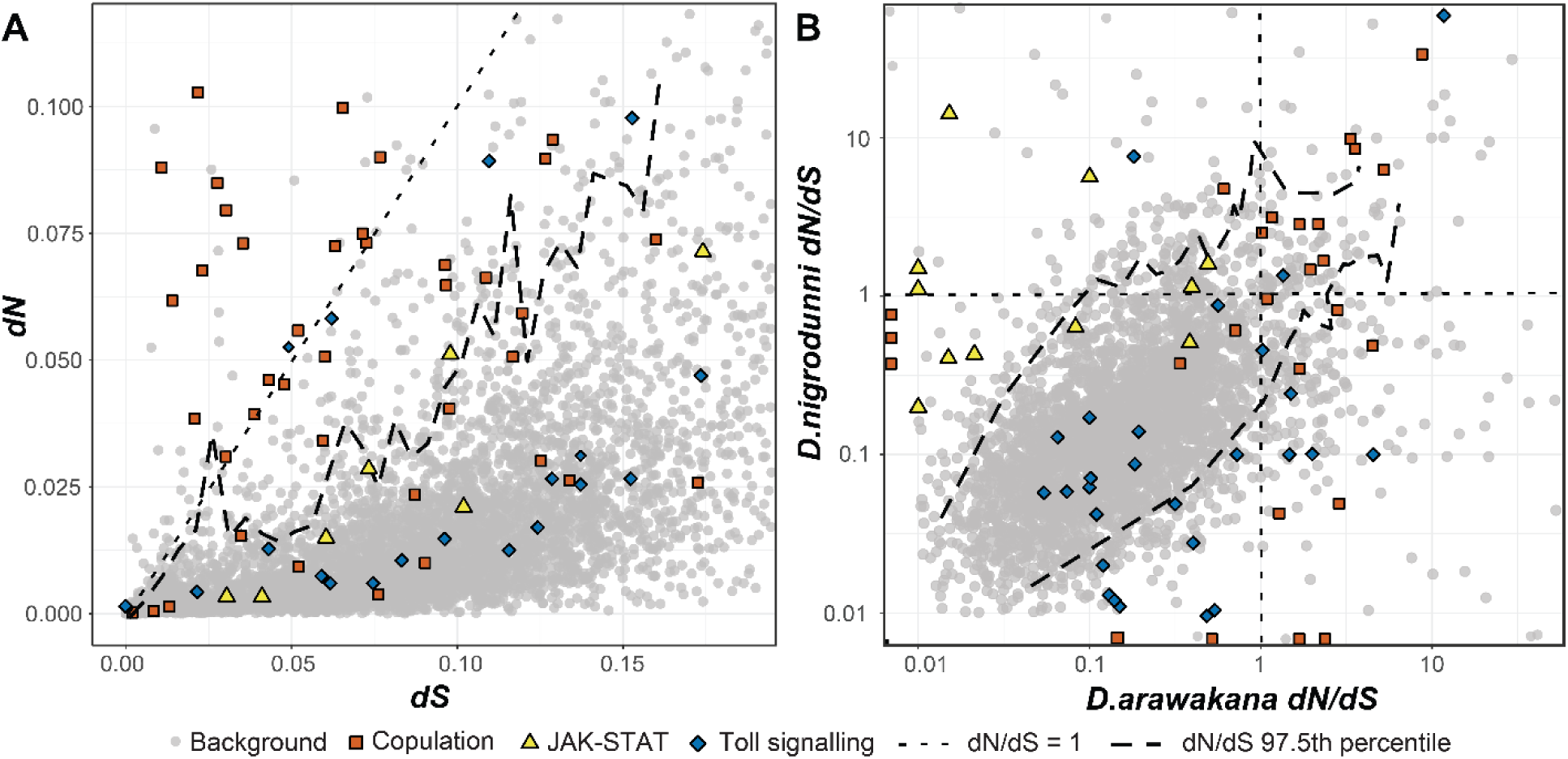
Rates of evolution across the *Drosophila dunni* phylogeny, showing non-synonymous divergence versus synonymous divergence across **A**. the whole phylogeny and **B**. comparing the proportion of non-synonymous to synonymous divergence between *D. nigrodunni* and *D. arawakana*. JAK-STAT, Toll and seminal fluid proteins are highlighted due to their enrichments in one or the other species.

When examining the repetitive content of each species, we see an expansion of Helitrons and LTRs along the *D. dunni/D. similis* branch, resulting in higher TE content in these two species compared to *D. nigrodunni/D. arawakana* (Supplementary Figure 5). We also find species-specific expansions of satellites, particularly in *D. arawakana* and *D. nigrodunni*, where ∼4% of the genome appears to be satellite sequences exclusive to that species (Supplementary Figure 5).

**Figure 5:**
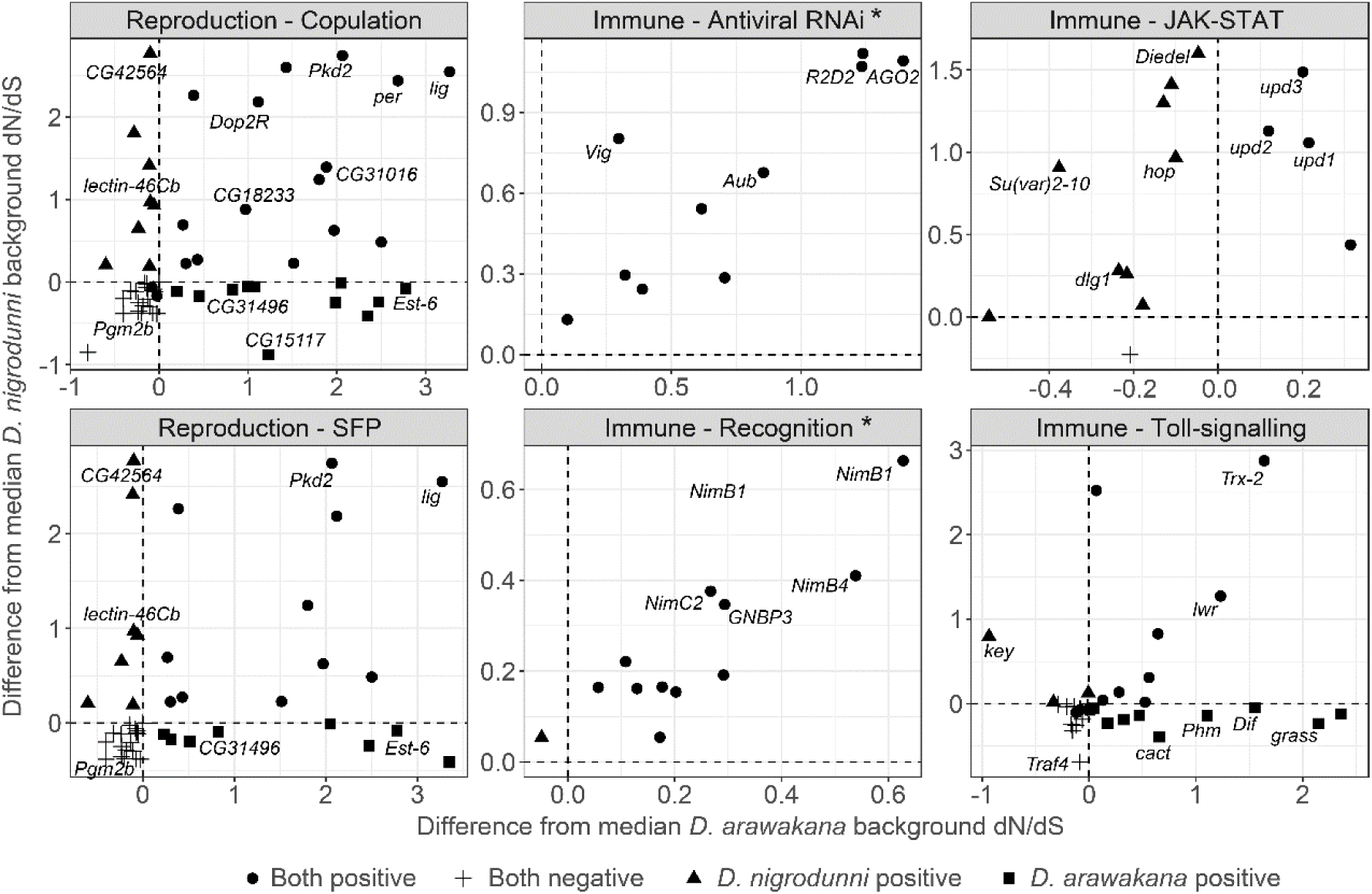
Difference of dN/dS between focal genes in specific functional categories and their nearby background genes. We find different insemination proteins and seminal fluid proteins are rapidly evolving between *D. nigrodunni* and *D. arawakana*. A selection of genes in each category are labelled by name in each plot. Plots are labelled with a * if we find a positive correlation between the two axes (*p*-value < 0.05).

We identified orthologous genes across species using BLAST (ALTSCHUL *et al*. 1990) with *D. innubila* as an outgroup when possible. For each group of orthologous genes, we identified the proportion of synonymous (dS) substitutions and amino acid changing, nonsynonymous substitutions (dN) (per possible synonymous or nonsynonymous substitution, respectively) occurring on each branch of the phylogeny using codeML (branch-based approach, model 0) (Yang 2007). We also estimated these substitution rates across the entire *dunni* group phylogeny (sites-based approach, model 7 & 8) (Yang 2007). This allowed us to calculate dN/dS to identify genes showing signatures of rapid or unconstrainted evolution on any branch of the phylogeny, or across the entire tree. For the dN/dS estimates on each branch, we identified genes in the upper 97.5^th^ percentile for dN/dS in windows of 0.01 dS. dN/dS in *D. nigrodunni* is significantly correlated with dN/dS in *D. arawakana* (Figure 2B), as well as in all other pairwise species comparisons (Supplementary Table 6, Pearson’s correlation coefficient = 0.844, t = 7.3774, df = 7569, *p*-value = 1.786e-13), and that similar proteins are rapidly evolving across the entire group. Copulation proteins (specifically seminal fluid proteins) are overrepresented among the most rapidly evolving genes on every branch of the *dunni* group phylogeny (Supplementary Table 6, *p*-value < 0.05 after multiple testing correction). This is consistent with rapid evolution occurring in genes involved in the reproductive conflict between the sexes (Figure 4) (Haerty *et al*. 2007). While not significant outliers, we also find that immune recognition proteins, antiviral RNA and piRNA pathways are also rapidly evolving in some species, consistent with arms races between the species and their parasites (Supplementary Table 6).

Rapidly evolving genes may provide clues into the selective forces acting on species since their divergence. For the main species pairs of interest (e.g. *D. nigrodunni* and *D. arawakana*) we identified genes in the upper 97.5^th^ percentile for windows of dN/dS in the other species, to find genes rapidly evolving in one species but not the other (Figure 2B). As expected, copulation-associated proteins were in the upper 97.5^th^ percentile for both species, while genes in the Toll immune pathway are rapidly evolving in *D arawakana* but not *D. nigrodunni*, conversely the JAK-STAT immune pathway is rapidly evolving in *D. nigrodunni* but not *D. arawakana* (Supplementary Table 6, Figure 4B). These results suggest each species may differ in their primary pathogen, resulting in context dependent immune evolution, as seen elsewhere in the *Drosophila* subgenus (Obbard *et al*. 2009; Hill *et al*. 2019).

We sought to confirm the rapid evolution of reproductive pathways and immune pathways after controlling for the background rate of evolution. We found the difference between dN/dS for each immune and reproductive gene and genes at neighboring loci on the chromosome (within 100kbp), of similar levels of divergence (+-0.01 dS). We find significantly elevated rates of evolution of antiviral genes, insemination genes and seminal fluid proteins across the entire phylogeny (Figure 5, one-sided T-test mu = 0, *p*-value = 0.0434). We also find a significant correlation between differences in *D. arawakana* and *D. nigrodunni* for antiviral genes (Pearson’s correlation = 0.795, t-value = 2.163, *p*-value = 0.0288), immune recognition genes (Pearson’s correlation = 0.877, t-value = 5.791, *p*-value = 0.000175) and piRNA genes (Pearson’s correlation = 0.659, t-value = 3.506, *p*-value = 0.00292). The highest average rate of evolution occurred seminal fluid proteins on the *D. nigrodunni* and *D. arawakana* branches (Figure 5, one-sided T-test, mu = 0, *p*-value < 0.05). Consistent with previous results we find elevated rates of evolution of the Toll signaling pathway in *D. arawakana*, and JAK-STAT in *D. nigrodunni*. Interestingly, when comparing the specific genes rapidly evolving between *D. nigrodunni* and *D. arawakana*, the specific insemination and seminal fluid genes are mostly evolving at different rates between species (Figure 5), while the other rapidly evolving genes are consistent between species (Figure 4B). Consistent with this, we find no correlation between measures between *D. arawakana* and *D. nigrodunni* in copulation (Pearson’s correlation = 0.187, t-value = 1.417, *p*-value = 0.162), seminal fluid proteins (Pearson’s correlation = 0.0341, t-value = 0.224, *p*-value = 0.823), JAK-STAT (Pearson’s correlation = 0.185, t-value = 0.625, *p*-value = 0.545) or Toll-signaling proteins (Pearson’s correlation = 0.450, t-value = 1.334, *p*-value = 0.224). This could suggest a difference in importance of insemination proteins between the species and could even suggest a functional divergence (Haerty *et al*. 2007).

Using orthology to *D. innubila*, we also identified duplications relative to these two species in each *dunni* group genome, and specific to each species. Consistent with the estimates in rates of evolution, we find enrichments of duplications in cell motility and copulation across the entire phylogeny (Supplementary Figure 6, Supplementary Table 7). We also find enrichments of duplications in Toll signaling genes in *D. arawakana* (*p*-value = 0.000569, enrichment = 5.44). Overall this suggests that the pathways showing elevated levels of nucleotide divergence (namely Toll and Copulation genes) also have more copy number variation between species than expected.

## Discussion

*Drosophila* species have served as prominent models in genetics research, including in understanding the divergence between populations and the evolution of species. This is facilitated by the extensive genetic tools available in the species group to identify the genetic basis of reproductive isolation, both prezygotic and postzygotic. Many islands contain endemic species of *Drosophila* with differing levels of isolation. For example, the island endemics in the *Drosophila simulans* complex (Cabot *et al*. 1994; Kliman *et al*. 2000; Matute and Ayroles 2014), with *D. mauritiana, D. simulans* and *D. sechellia* have served as a rich system for understanding reproductive isolation (Cabot *et al*. 1994; Kliman *et al*. 2000). Like the *Drosophila simulans* complex, the *Drosophila dunni* species subgroup has radiated across a chain of islands (Heed 1962), though with easier to define species relationships than is seen in the *simulans* subcomplex (Cabot *et al*. 1994; Kliman *et al*. 2000; Matute *et al*. 2014). Due to the recent radiation of this group, many species pairs in the *dunni* subgroup produce offspring (Stalker and Streisinger 1953; Heed 1962), some of which are fertile, and so provide a potentially useful model system for dissecting the genetics of reproductive isolation.

Here, we assessed the extent of hybrid incompatibilities between species of the *dunni* subgroup, focusing on post-mating-prezygotic incompatibilities. We then sequenced and assembled the species genomes to identify highly divergent and rapidly evolving genes. Between *D. nigrodunni* and *D. arawakana*, we find elevated divergence of several immune system pathways, as well as divergence in genes involved in copulation. This divergence fits with the hybrid male inviability between these two species, as well as the reduced survival of females following insemination by a heterospecific male. Consistent with the divergence in the seminal fluid proteins, we find evidence of an insemination reaction-like swelling of the reproductive tract (Knowles and Markow 2001), and a decrease in hybrid mating compared to within species.

Strangely, in this study most of the striking differences appear when comparing *D. nigrodunni* and *D. arawakana* (Figures 1-5). This pair is slightly less diverged than other pairings within the group (Supplementary Figure 1) and are allopatrically separated (Heed 1962; Wilder and Hollocher 2003), allowing for the neutral accumulation of substitutions with a reduced chance of introgression or reinforcement (Coyne and Orr 1989; Coyne and Orr 2004). Due to this reduced divergence and reduced incidence of incompatibilities (Orr 1995; Welch 2004), we may have caught this species pair at the opportune time where these hybrid incompatible effects are visible, while other species pairs are too far diverged (Figure 1).

The functional annotation of the more diverged genes may also provide us with clues as to how these species are diverging. As we find premating-behavior proteins are divergent between *D. arawakana* and *D. nigrodunni*, this may result in a divergence in premating behavior, resulting in the reduced rate of hybrid matings scored (Figure 3). We also see no difference in the proportion of hybrid matings after 24 hours and 48 hours, suggesting that in these cases, if a female has rejected all males, she may not change her mind later (Coyne and Orr 2004; GourbiÈre and Mallet 2010; Turissini *et al*. 2017). Hybridization between island-endemic flies separated by ∼500 kilometers of ocean may be unlikely (Coyne *et al*. 1982), but selection against hybridization between our focal species and other *dunni* group species may have led to the evolution of reinforcement against heterospecific mating (GourbiÈre and Mallet 2010; Turissini *et al*. 2017). We also find seminal fluid and copulation proteins are rapidly diverging between species (Figure 2) and find an increased incidence of swollen and deformed reproductive tracts, consistent with an insemination reaction-like effect and a toxic incompatibility between the SFPs and their environment (Figures 2 and 3) (Markow and Ankney 1988; Knowles and Markow 2001). In fact, studies in other species have also identified post-mating-prezygotic incompatibilities to be a driver of isolation between species, even in cases with gene flow (Gavrilets and Waxman 2002; Gavrilets 2003; Larson *et al*. 2012; Larson *et al*. 2013; AHMED-BRAIMAH 2016; Turissini *et al*. 2017). A recent study identified the upregulation of the JAK-STAT pathway (a stress response pathway) in *Drosophila* females following heterospecific mating, likely due to the negative effects of the accessory gland proteins (AHMED-BRAIMAH *et al*. 2020). The rapid evolution of JAK-STAT proteins in *D. nigrodunni* could also be due to this species requiring a well-adapted stress response pathway, given its negative reaction to heterospecific matings (Figures 1-3).

Several of the functional gene categories identified in this study as highly divergent between species are also promising regions for future study, particularly when focusing on immune evolution. Our findings are consistent with other studies that find immune proteins are more rapidly evolving than background genes (Sackton *et al*. 2007; Obbard *et al*. 2009; Shultz and Sackton 2019), consistent with an arms-race between the host and its pathogens. However, in the species studied here, we find several cases of species-specific rapid evolution of an immune pathway, such as the rapid evolution of JAK-STAT in *D. nigrodunni* (Figures 4 and 5). As immune pathways are constantly evolving in response to their pathogens, this could be explained by differences in immune pathogens in this species group (Sackton *et al*. 2007; Unckless *et al*. 2016; Hill *et al*. 2019). Hypothetically, the lack of any substantive natural Gram-Negative bacterial pathogens in *D. dunni* would result in a lack of divergence in the IMD pathway, the immune pathway associated with the resisting Gram-Negative bacteria. While a lack of fungal or Gram-Positive bacterial pathogens in *D. nigrodunni* could result in the lack of evolution of the Toll pathway, but rampant evolution in *D. arawakana* (Figures 4 and 5).

The repetitive content also appears to be diverging rapidly across this species complex (Supplementary Figure 5). This is commonly seen between species, given the elevated mutation rate/transposition of selfish factors compared to the rest of the genome (Kofler *et al*. 2012; Kofler *et al*. 2015; Adrion *et al*. 2019), and has been implicated in the formation of hybrid incompatibilities for several species (Satyaki *et al*. 2014). Consistent with this we find several TE families unique to specific species in the *dunni* complex. However, we did not find a significant excess of dysgenic ovaries in hybrid females compared to normal females (Fisher’s exact text *p*-value > 0.05 for all cases). Several cases of hybrid incompatibilities caused by differences in TE content results in sterility caused by maternally inherited factors over paternally inherited (as is usually seen). This may be due to the absence of maternally loaded silencing RNAs against specific TEs (Bingham *et al*. 1982; Aravin *et al*. 2007; Brennecke *et al*. 2008). If this were the case, we would expect the hybrid sterility to be in the opposite direction to what we observe, with sterile females (Figure 1, Supplementary Figure 5) (Kidwell *et al*. 1977), and so do not expect the hybrid incompatibilities seen here to be caused by repetitive content. However, this is a simplistic view of the effects of transposon activity on hybrid fertility, given the complex hybrid dysgenesis cases seen in *D. virilis* (Petrov et al. 1995; EVGEN’EV et al. 1997; Erwin et al. 2015), and even the complex cases of tolerance to dysgenesis seen in the supposedly simple case in *D. melanogaster* (Kelleher et al. 2018), so may require further study to fully understand if TEs play a role in the divergence of the *dunni* complex.

Overall, our findings suggest that the rapid divergence of reproductive genes has led to incompatibilities between species in the *dunni* group, including inviable male offspring and the insemination reaction associated with reduced female survival. We also find multiple areas for further investigation in the *D. dunni* group, either in immune evolution of continuing to investigate the speciation in this species group, suggesting promise in the future of research for this group.

## Supplementary Information

**Supplementary Table 1:** Table of next-generation sequencing information used in this survey with number of reads per sample and accession numbers per sample.

**Supplementary Table 2:** Average number of offspring from each set of crosses, either heterospecific or conspecific crosses after 1 week of mating and 1 week of egg laying. Table also shows the sex ratio of offspring and if offspring of each sex are fertile.

**Supplementary Table 3:** Table showing average synonymous divergence between each species pair.

**Supplementary Table 4:** Table summarizing genome assembly statistics of each species sequenced and assembled chromosomes, including number of scaffolds for each chromosome, the length of each chromosome, and coding and intronic proportions.

**Supplementary Table 5:** Summary statistics of genomes sequenced assembled and annotated in this study, including number of genes, the number of these that have orthologs in *D. virilis* and *D. melanogaster*, as well as statistics regarding size of these genes.

**Supplementary Table 6:** Gene categories enriched for high dN/dS (either the upper 95^th^ percentile or dN/dS > 1) across the entire phylogeny and on each species branch, also the upper outliers for *D. nigrodunni* and *D. arawakana* relative to the other species.

**Supplementary Table 7:** Gene categories enriched for duplications on each branch of the *D. dunni* species phylogeny, relative to unduplicated genes.

**Supplementary Table 8:** dN/dS statistics calculated using codeML for the entire *dunni* phylogeny and on each branch.

**Supplementary Figure 1:**
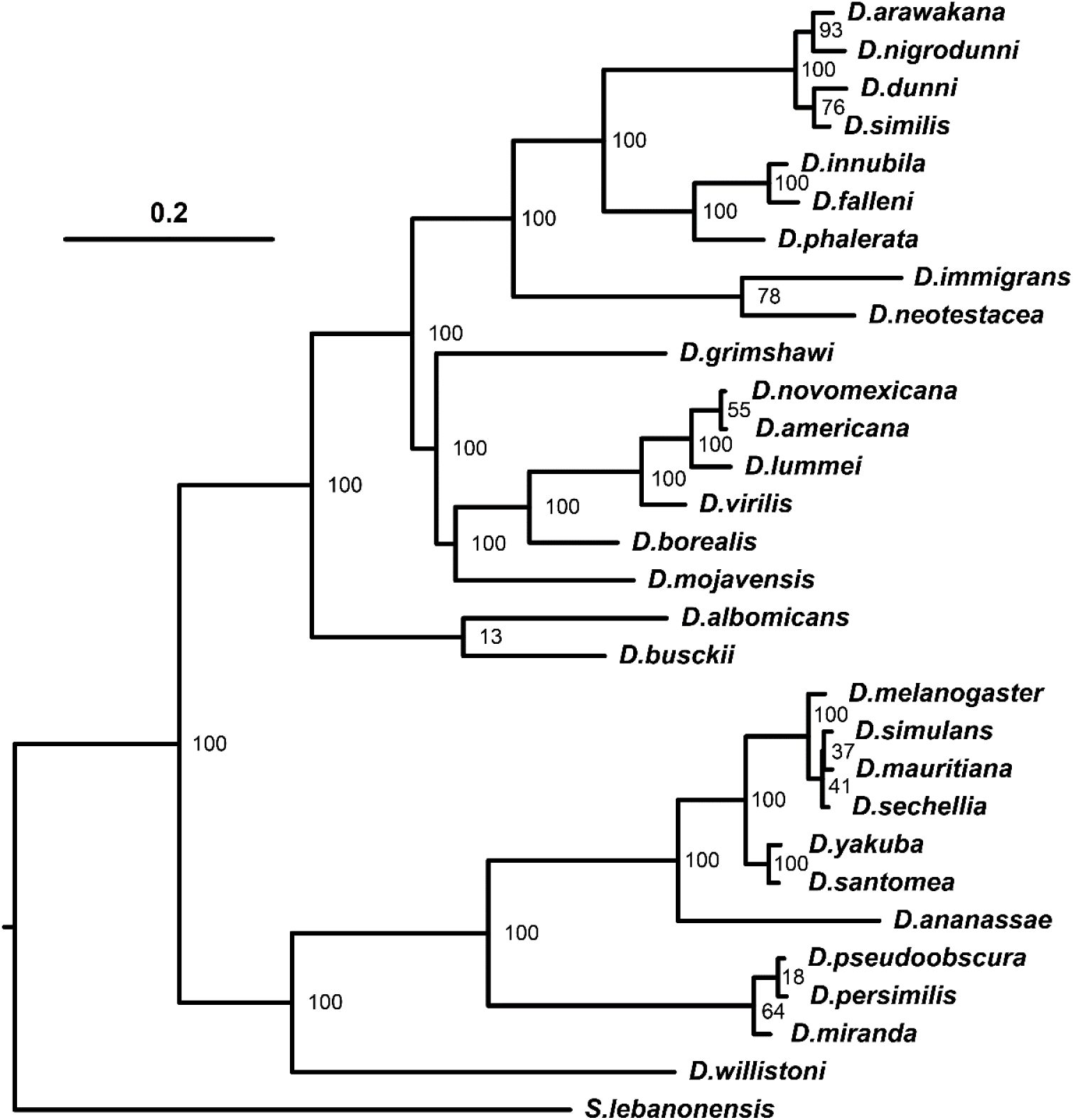
Phylogeny of the *dunni* group relative to other *Drosophila* species. Phylogeny was calculated using PhyML (GUINDON *et al*. 2010), finding the consensus of 100 genes, with bootstrap values (the number that match out of 100) shown at nodes.

**Supplementary Figure 2:**
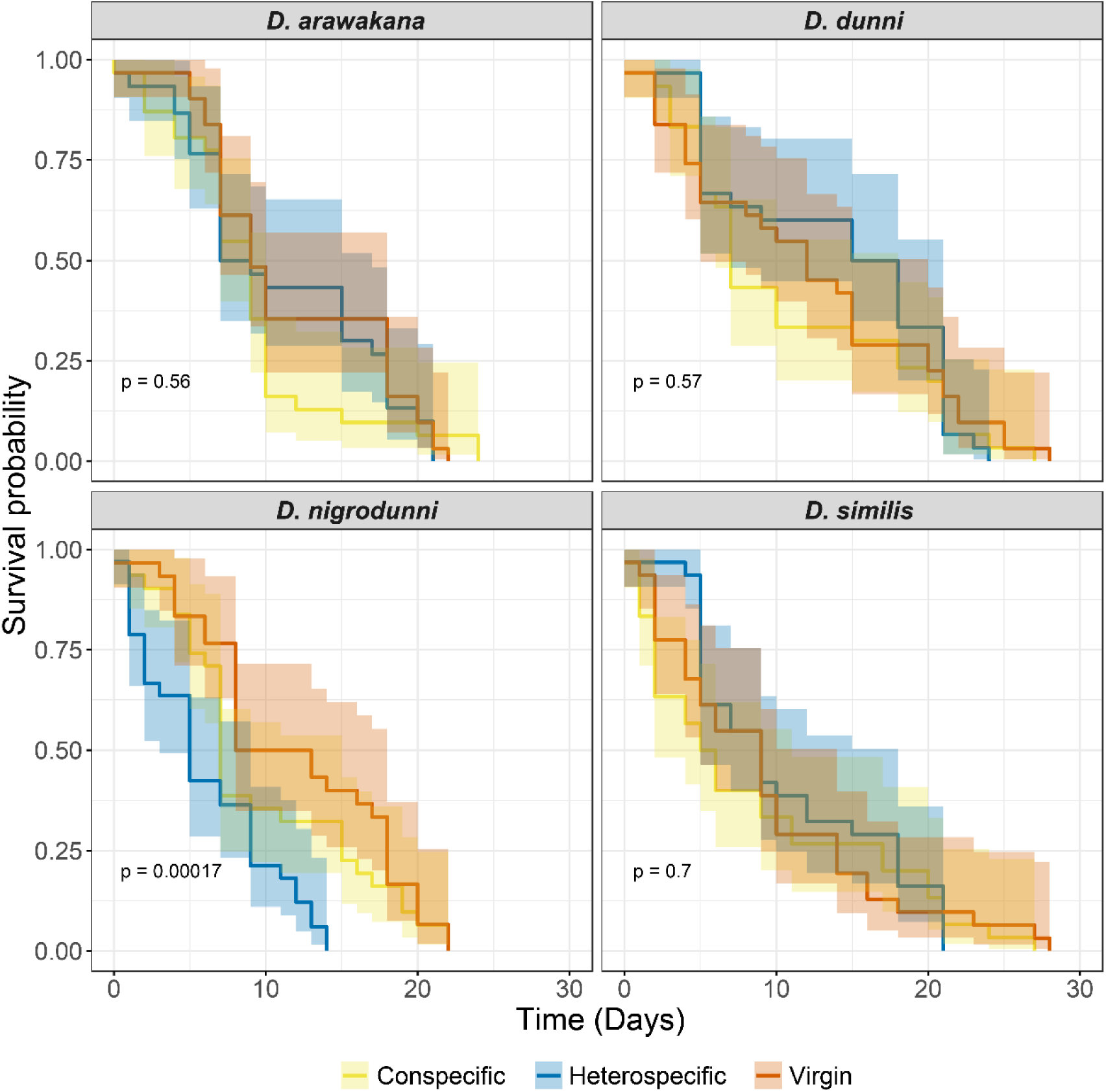
Survival probability of females for each species used in each cross, compared to virgin female survival. Crosses following curing of the strain with Tetracycline-Hydrochloride. Females are separated by species, and grouped as virgins, conspecific crossed (crossed to own species), heterospecific crossed (crossed to a different species). In the case of heterospecific crosses, *D. arawakana* is only crossed to *D. nigrodunni* and *D. dunni* is only crossed to *D. similis*.

**Supplementary Figure 3:**
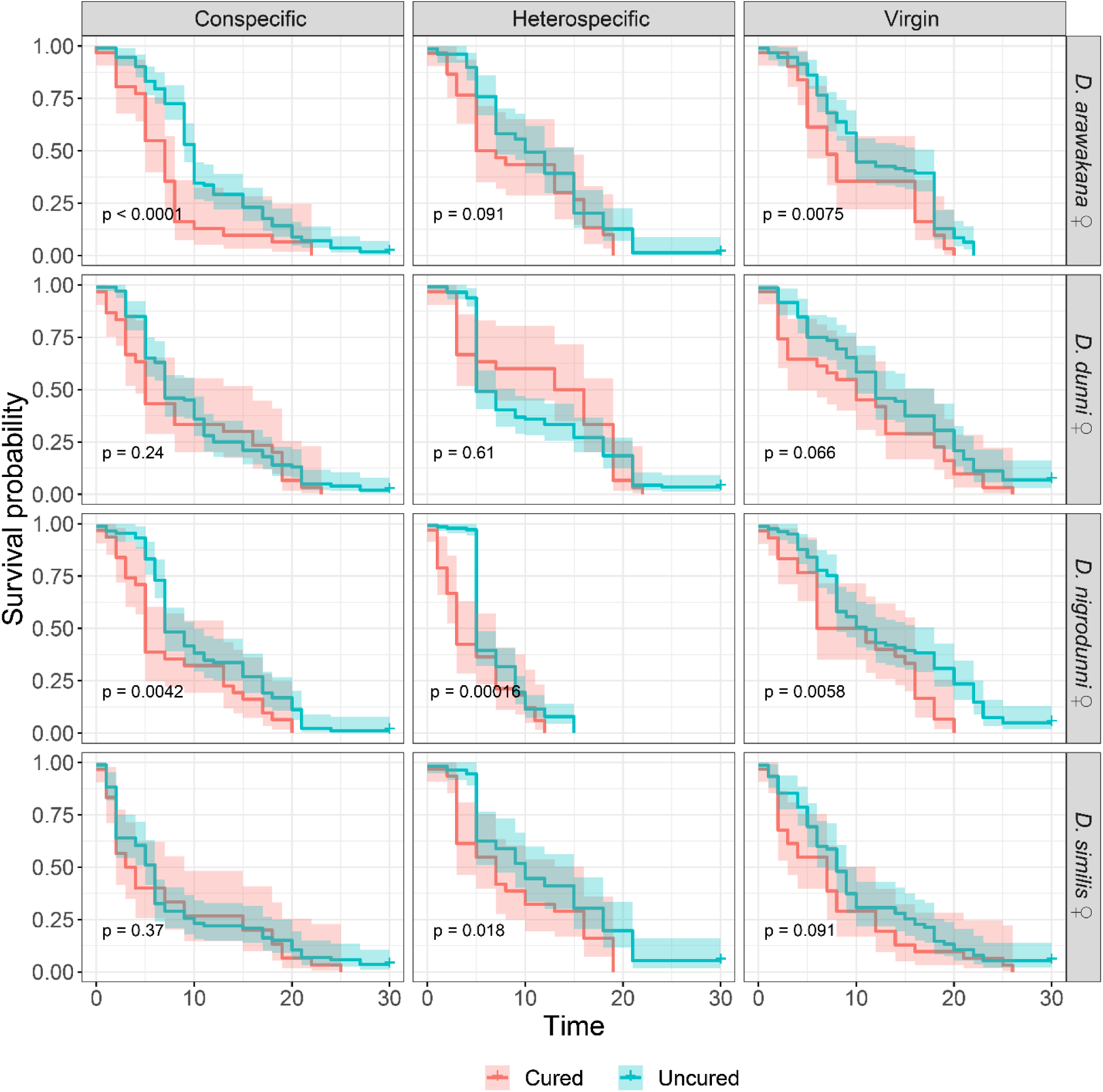
Difference in survival for different sets of crosses, comparing between survival of females before and after curing with Tetracycline-Hydrochloride.

**Supplementary Figure 4:**
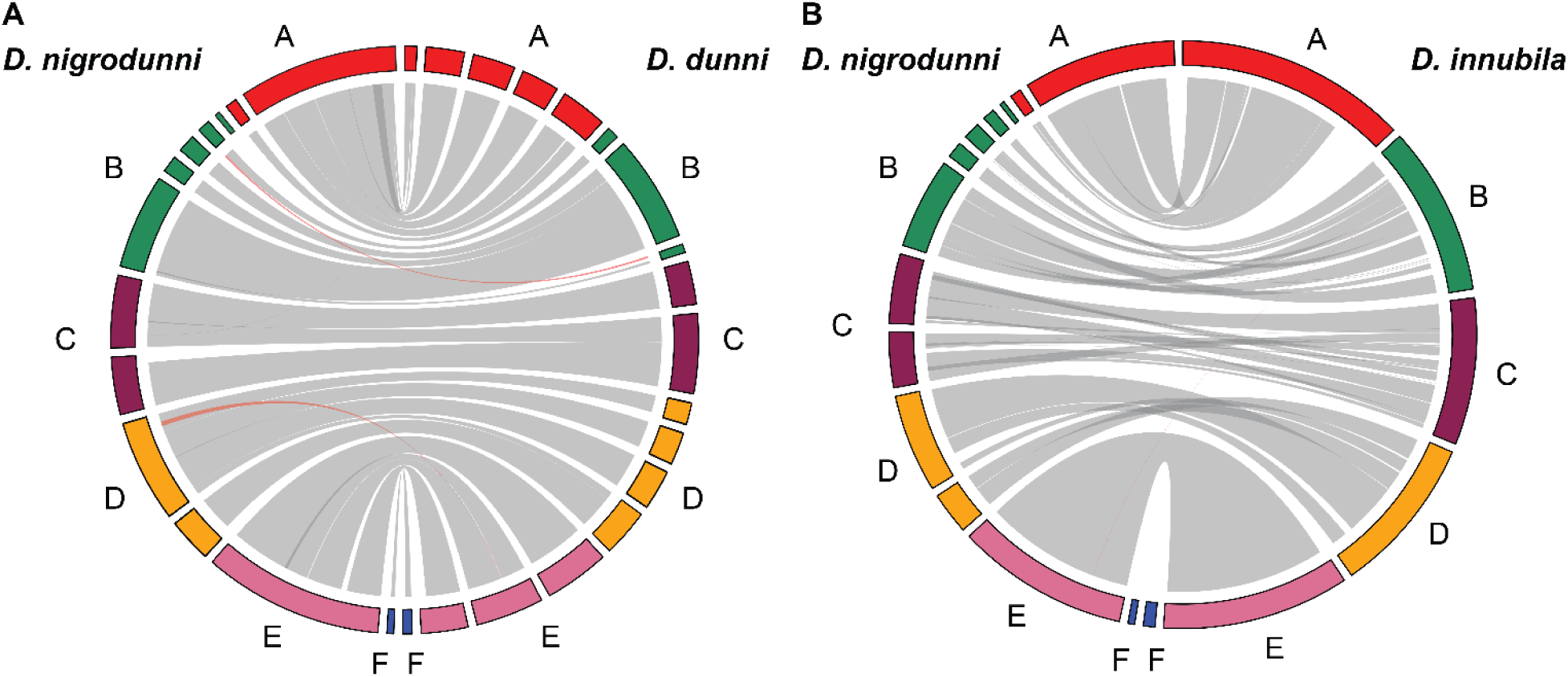
Orthologous regions between the *D. nigrodunni* genome, *D. dunni* genome and *D. innubila* genome. Syntenic regions on the same chromosome (shown as Muller elements, A-F) are labelled with grey ribbons, while syntenic regions between difference chromosomes are labelled in red.

**Supplementary Figure 5:**
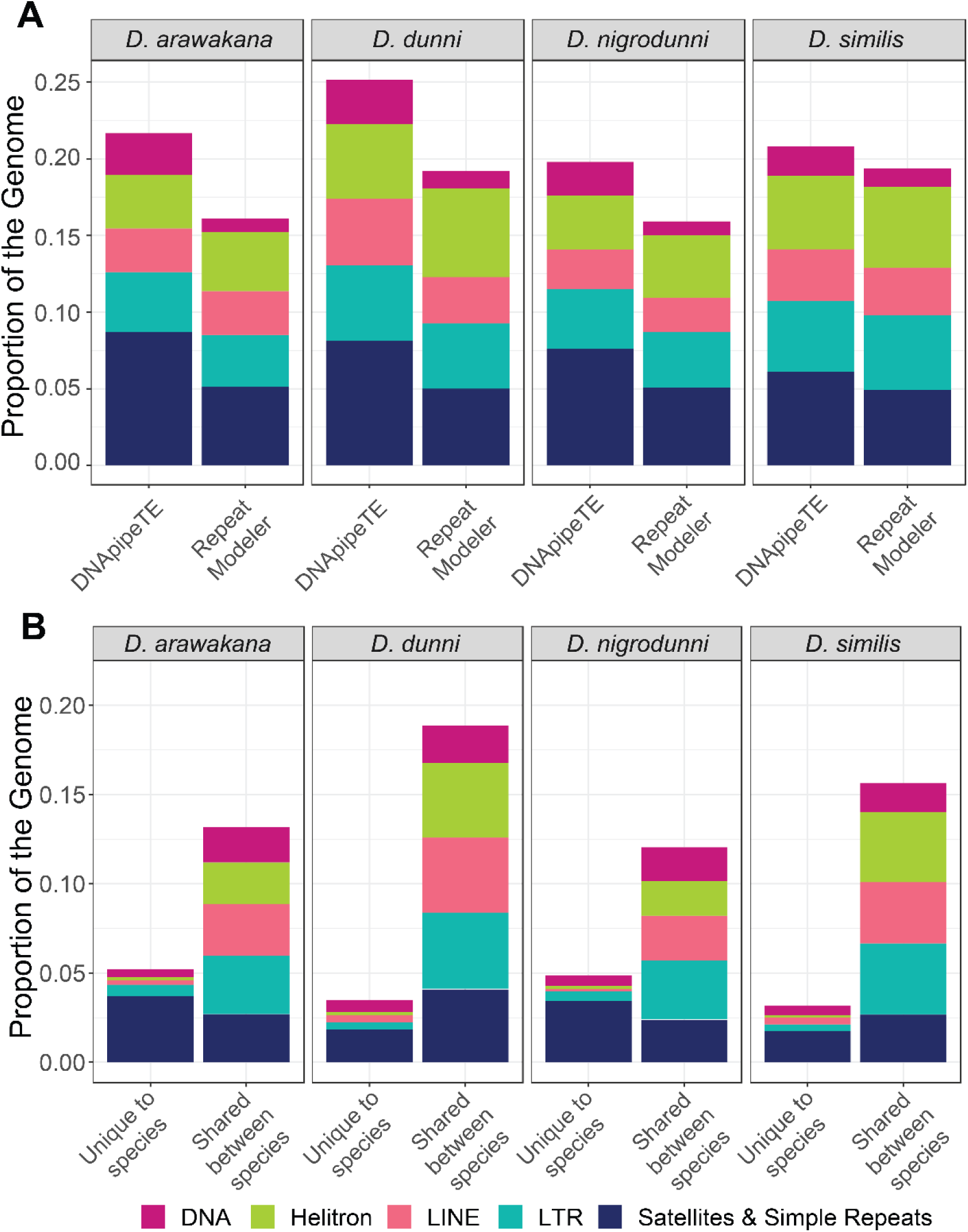
Proportion of each genome made up of repetitive content (colored by classification of repetitive content) For simplicity, the satellite category contains Satellites, microsatellites, simple repeats tandem repeats and low complexity regions. **A**. Comparison of TE annotation between two tools, DNApipeTE and Repeatmodeler. **B**. Comparison of TE content across species and if that content is shared between species or is unique to one species.

**Supplementary Figure 6:**
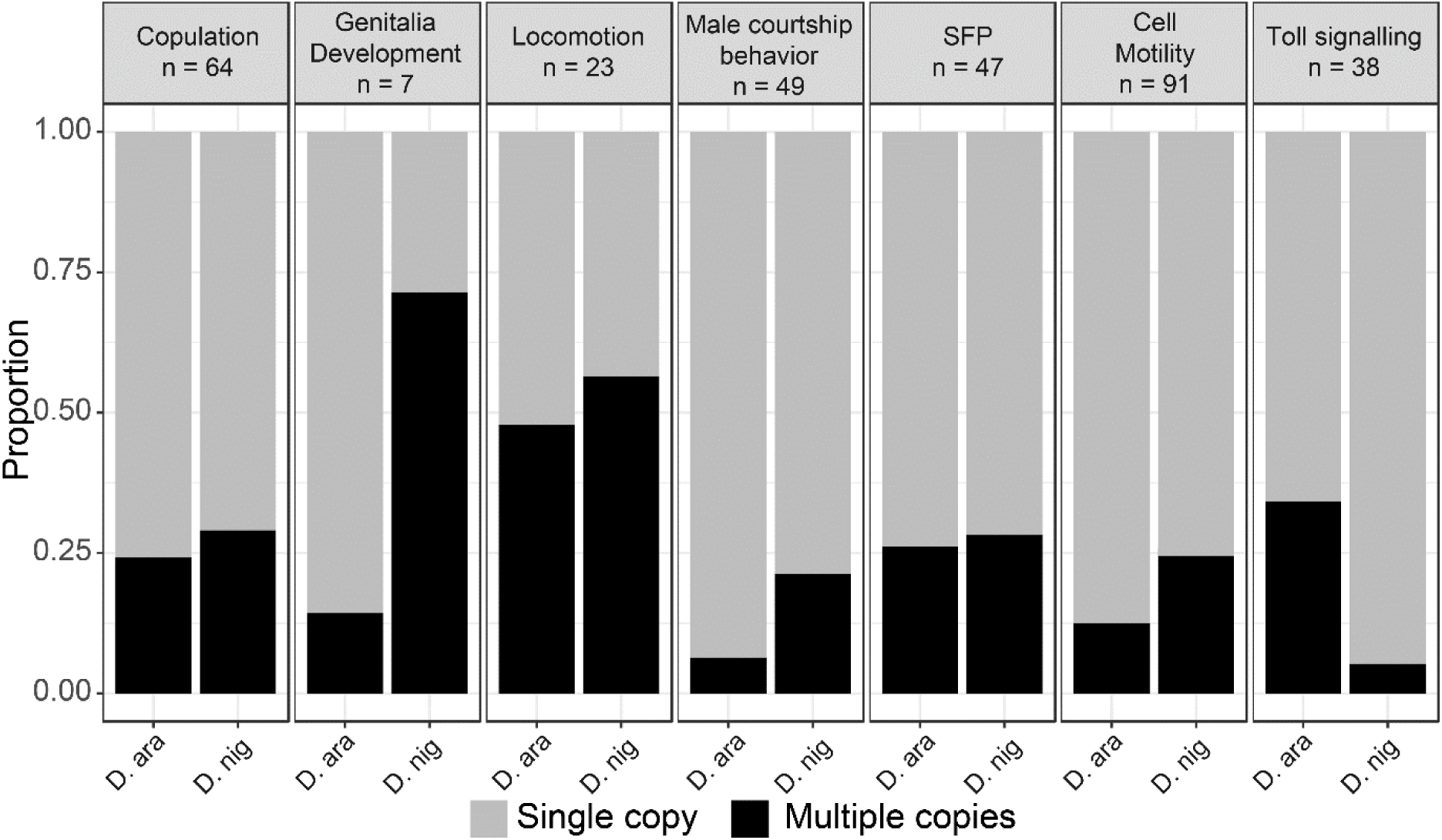
Proportion of gene categories with enrichments of duplications in *D. nigrodunni* (D. nig), *D. arawakana* (D. ara) or both species. N = number of genes per category.

## References

Adrion, J. R., D. J. Begun and M. W. Hahn, 2019 Patterns of transposable element variation and clinality in *Drosophila*. Mol Ecol 28: 1523–1536.

Ahmed-Braimah, Y., 2016 Multiple Genes Cause Postmating Prezygotic Reproductive Isolation in the *Drosophila virilis* Group. G3: Genes|Genomes|Genetics 6: 4067–4076.

Ahmed-Braimah, Y. H., M. F. Wolfner and A. G. Clark, 2020 Differences in post-mating transcriptional responses between conspecific and heterospecific matings in *Drosophila*. Biorxiv.

Altschul, S. F., W. Gish, W. Miller, E. W. Myers and D. J. Lipman, 1990 Basic local alignment search tool. Journal of Molecular Biology 215: 403–410.

Andres, J. A., L. S. Maroja and R. G. Harrison, 2008 Searching for candidate speciation genes using a proteomic approach: seminal proteins in field crickets. Proc Biol Sci 275: 1975–1983.

Aravin, A. A., G. J. Hannon and J. Brennecke, 2007 The piwi-piRNA pathway provides an adaptive defense in the transposon arms race. Science 318: 761–764.

Bingham, P. M., M. G. Kidwell and G. M. Rubin, 1982 The molecular basis of P-M hybrid dysgenesis: the role of the *P*-element, a P-strain-specific transposon family. Cell 29: 995–1004.

Brekke, T. D., and J. M. Good, 2014 Parent-of-origin growth effects and the evolution of hybrid inviability in dwarf hamsters. Evolution 68: 3134–3148.

Brennecke, J., C. D. Malone, A. A. Aravin, R. Sachidanandam, A. Stark et al., 2008 An epigenetic role for maternally inherited piRNAs in transposon silencing. Science 322: 1387–1392.

Broad-Institute, 2017 Picard, pp., http://broadinstitute.github.io/picard.

Buffalo, V., 2018 Scythe.

Cabot, L., A. W. Davis, N. A. Johnson and C.-i. Wu, 1994 Genetics of reproductive isolation in the *Drosophila simulans* clade: complex epistasis underlying hybrid male sterility. Genetics 137: 175–189.

Cenzi De Ré, F., E. L. S. Loreto and L. J. Robe, 2010 Gene and species trees reveal mitochondrial and nuclear discordance in the *Drosophila cardini* group (Diptera: Drosophilidae). Invertebrate Biology 129: 353–367.

Chapman, T., J. Hutchings and L. Partridge, 1993 No reduction in the cost of m ating for *Drosophila melanogaster* females mating w ith spermless males. Proceedings R. Soc. 253: 211–217.

Cordeiro, J., D. C. De Toni, G. de Souza da Silva and V. L. da Silva Valente, 2014 Chromosomal evolution in the *Drosophila cardini* group (Diptera: Drosophilidae): photomaps and inversion analysis. Genetica 142: 461–472.

Coyne, J., and H. A. Orr, 2004 Speciation.

Coyne, J. A., I. A. Boussy, T. Prout, S. H. Bryant, J. S. Jones et al., 1982 Long Distance Migration of *Drosophila*. The American Naturalist 11: 589–595.

Coyne, J. A., and H. A. Orr, 1989 Patterns of speciation in *Drosophila.* Evolution 43: 362–381.

Darling, A. C. E., B. Mau, F. R. Blattner and N. T. Perna, 2004 Mauve : Multiple Alignment of Conserved Genomic Sequence With Rearrangements Mauve : Multiple Alignment of Conserved Genomic Sequence With Rearrangements. Genome Res 14: 1394–1403.

DePristo, M. A., E. Banks, R. Poplin, K. V. Garimella, J. R. Maguire et al., 2011 A framework for variation discovery and genotyping using next-generation DNA sequencing data. Nature genetics 43: 491–498.

Eden, E., R. Navon, I. Steinfeld, D. Lipson and Z. Yakhini, 2009 GOrilla: A tool for discovery and visualization of enriched GO terms in ranked gene lists. BMC Bioinformatics 10: 1–7.

Erwin, A. A., M. A. Galdos, M. L. Wickersheim, C. C. Harrison, K. D. Marr et al., 2015 piRNAs Are Associated with Diverse Transgenerational Effects on Gene and Transposon Expression in a Hybrid Dysgenic Syndrome of D. virilis. PLoS Genetics 11: 1–33.

Evgen’ev, M. B., H. Zelentsova, N. Shostak, M. Kozitsina, V. Barskyi et al., 1997 Penelope, a new family of transposable elements and its possible role in hybrid dysgenesis in Drosophila virilis. Proceedings of the National Academy of Sciences of the United States of America 94: 196–201.

Gavrilets, S., 2003 Perspective: models of speciation: what have we learned in 40 years? Evolution; international journal of organic evolution 57: 2197–2215.

Gavrilets, S., and D. Waxman, 2002 Sympatric speciation by sexual conflict. PNAS 16: 10533–10538.

Glor, R. E., J. B. Losos and A. Larson, 2005 Out of Cuba: overwater dispersal and speciation among lizards in the Anolis carolinensis subgroup. Mol Ecol 14: 2419–2432.

Goubert, C., L. Modolo, C. Vieira, C. V. Moro, P. Mavingui et al., 2015 De novo assembly and annotation of the Asian tiger mosquito (*Aedes albopictus*) repeatome with dnaPipeTE from raw genomic reads and comparative analysis with the yellow fever mosquito (*Aedes aegypti*). Genome Biology and Evolution 7: 1192–1205.

Gourbière, S., and J. Mallet, 2010 Are species real? The shape of the species boundary with exponential failure, reinforcement, and the “missing snowball”. Evolution; international journal of organic evolution 64: 1–24.

Gramates, L. S., S. J. Marygold, G. Dos Santos, J. M. Urbano, G. Antonazzo et al., 2017 FlyBase at 25: Looking to the future. Nucleic Acids Research 45: D663–D671.

Grant, B., 1983 On the Relationship Between Average Copulation Duration and Insemination Reaction in the Genus Drosophila. Evolution 37: 854–856.

Guindon, S., J.-F. Dufayard, V. Lefort, M. Anisimova, W. Hordijk et al., 2010 New algorithms and methods to estimate maximum-likelihood phylogenies: assessing the performance of PhyML 3.0. Systematic biology 59: 307–321.

Haerty, W., S. Jagadeeshan, R. J. Kulathinal, A. Wong, K. Ravi Ram et al., 2007 Evolution in the fast lane: rapidly evolving sex-related genes in *Drosophila*. Genetics 177: 1321–1335.

Haldane, J., 1922 Sex ratio and unisexual sterility in animal hybrids. J Genet 12: 101–109.

Hamilton, P. T., J. S. Leong, B. F. Koop and S. J. Perlman, 2014 Transcriptional responses in a *Drosophila* defensive symbiosis. Molecular Ecology 23: 1558–1570.

Heed, W. B., 1962 IX. Genetic characteristics of Island Populations, pp. 173–206 in Univ Texas Publ.

Hill, T., B. Koseva and R. L. Unckless, 2019 The genome of *Drosophila innubila* reveals lineage-specific patterns of selection in immune genes. Molecular Biology and Evolution: 1–36.

Hill, T., and R. L. Unckless, 2019 A Deep Learning Approach for Detecting Copy Number Variation in Next-Generation Sequencing Data. G3 9.

Hollocher, H., J. L. Hatcher and E. G. Dyreson, 2000 Evolution of abdominal pigmentation differences across species in the *Drosophila dunni* subgroup. Evolution 54: 2046–2056.

Holt, C., and M. Yandell, 2011 MAKER2 : an annotation pipeline and genome-database management tool for second-generation genome projects. BMC Bioinformatics 12: 491.

Howard, D. J., J. L. Marshall, D. D. Hampton, S. C. Britch, M. L. Draney et al., 2002 The genetics of reproductive isolation: a retrospective and prospective look with comments on ground crickets. The American naturalist 159 Suppl: S8–S21.

Jain, M., H. E. Olsen, B. Paten and M. Akeson, 2016 The Oxford Nanopore MinION: Delivery of nanopore sequencing to the genomics community. Genome Biology 17: 1–11.

Johnson, A. D., R. E. Handsaker, S. L. Pulit, M. M. Nizzari, C. J. O’Donnell et al., 2008 SNAP: a web-based tool for identification and annotation of proxy SNPs using HapMap. Bioinformatics 24: 2938–2939.

Joshi, N., and J. Fass, 2011 Sickle: A sliding window, adaptive, quality-based trimming tool for fastQ files. 1.33.

Kassambara, A., M. Kosinski and P. Biecek, 2017 survminer: Drawing Survival Curves using’ggplot2’. R package version 0.3 1.

Katoh, K., K. Misawa, K.-i. Kuma and T. Miyata, 2002 MAFFT: a novel method for rapid multiple sequence alignment based on fast Fourier transform. Nucleic acids research 30: 3059–3066.

Kelleher, E. S., J. Jaweria, U. Akoma, L. Ortega and W. Tang, 2018 QTL mapping of natural variation reveals that the developmental regulator bruno reduces tolerance to P-element transposition in the Drosophila female germline. PLoS Biol 16: e2006040.

Kidwell, J. F., M. G. Kidwell and J. A. Sved, 1977 Hybrid dysgenesis in *Drosophila melanogaster*: A syndrome of aberrant traits including mutation, sterility, and male recombination. Genetics 86: 813–833.

Kitano, J., J. A. Ross, S. Mori, M. Kume, F. C. Jones et al., 2009 A role for a neo-sex chromosome in stickleback speciation. Nature 461: 1079–1083.

Kitts, P. A., D. M. Church, F. Thibaud-Nissen, J. Choi, V. Hem et al., 2016 Assembly: a resource for assembled genomes at NCBI. Nucleic Acids Res 44: D73–80.

Kliman, R. M., P. Andolfatto, J. A. Coyne, F. Depaulis, M. Kreitman et al., 2000 The population genetics of the origin and divergence of the *Drosophila simulans* complex species. Genetics 156: 1913–1931.

Knowles, L. L., and T. A. Markow, 2001 Sexually antagonistic coevolution of a postmating-prezygotic reproductive character in desert *Drosophila*. PNAS 98.

Kofler, R., A. J. Betancourt and C. Schlötterer, 2012 Sequencing of pooled DNA Samples (Pool-Seq) uncovers complex dynamics of transposable element insertions in *Drosophila melanogaster*. PloS Genetics 8: 1–16.

Kofler, R., V. Nolte and C. Schlötterer, 2015 Tempo and mode of transposable element activity in *Drosophila*. PLoS Genet 11: e1005406.

Larson, E. L., J. A. Andres, S. M. Bogdanowicz and R. G. Harrison, 2013 Differential introgression in a mosaic hybrid zone reveals candidate barrier genes. Evolution 67: 3653–3661.

Larson, E. L., J. A. Andres and R. G. Harrison, 2012 Influence of the male ejaculate on post-mating prezygotic barriers in field crickets. PLoS One 7: e46202.

Le, S. Q., and O. Gascuel, 2008 An improved general amino acid replacement matrix. Molecular biology and evolution 25: 1307–1320.

Li, H., 2016 Minimap and miniasm: fast mapping and de novo assembly for noisy long sequences. Bioinformatics 32: 2103–2110.

Li, H., and R. Durbin, 2009 Fast and accurate short read alignment with Burrows-Wheeler transform. Bioinformatics (Oxford, England) 25: 1754–1760.

Li, H., B. Handsaker, A. Wysoker, T. Fennell, J. Ruan et al., 2009 The sequence alignment/map format and SAMtools. Bioinformatics (Oxford, England) 25: 2078–2079.

Liu, Y., and Y. Yang, 2013 Quick-merge sort algorithm based on multi-core linux, pp. 1578–1583 in Proceedings 2013 International Conference on Mechatronic Sciences, Electric Engineering and Computer (MEC). IEEE.

Löytynoja, A., 2014 Phylogeny-aware alignment with PRANK, pp. 155–170 in Multiple Sequence Alignment Methods, edited by D. J. Russell. Humana Press, Totowa, NJ.

Markow, T. A., and P. F. Ankney, 1988 Insemination Reaction in Drosophila: Found in Species Whose Males Contribute Material to Oocytes Before Fertilization. Evolution 42: 1097–1101.

Matute, D. R., and J. F. Ayroles, 2014 Hybridization occurs between Drosophila simulans and D. sechellia in the Seychelles archipelago. Journal of evolutionary biology 27: 1057–1068.

Matute, D. R., I. a. Butler, D. a. Turissini and J. A. Coyne, 2010 A test of the snowball theory for the rate of evolution of hybrid incompatibilities. Science (New York, N.Y.) 329: 1518–1521.

Matute, D. R., A. A. Comeault, E. Earley, A. Serrato-Capuchina, D. Peede et al., 2020 Rapid and Predictable Evolution of Admixed Populations Between Two <em>Drosophila</em> Species Pairs. Genetics 214: 211–230.

Matute, D. R., J. Gavin-Smyth and G. Liu, 2014 Variable post-zygotic isolation in *Drosophila melanogaster*/*D. simulans* hybrids. Journal of evolutionary biology 27: 1691–1705.

McKenna, A., M. Hanna, E. Banks, A. Sivachenko, K. Cibulskis et al., 2010 The Genome Analysis Toolkit: A MapReduce framework for analyzing next-generation DNA sequencing data. Proceedings of the International Conference on Intellectual Capital, Knowledge Management & Organizational Learning 20: 1297–1303.

McKinnon, J. S., and H. D. Rundle, 2002 Speciation in nature: the threespine stickleback model systems. Trends in Ecology & Evolution 17: 480 – 488.

Mendes, F., and M. W. Hahn, 2016 Gene tree discordance causes apparent substitution rate variation. Systematic Biology 65: 711–721.

Miller, W. J., L. Ehrman and D. Schneider, 2010 Infectious speciation revisited: Impact of symbiont-depletion on female fitness and mating behavior of *Drosophila paulistorum*. PLoS Pathogens 6: e1001214.

Moyle, L. C., and T. Nakazato, 2010 Hybrid incompatibility “snowballs” between Solanum species. Science (New York, N.Y.) 329: 1521–1523.

Narasimhan, V., P. Danecek, A. Scally, Y. Xue, C. Tyler-Smith et al., 2016 BCFtools/RoH: A hidden Markov model approach for detecting autozygosity from next-generation sequencing data. Bioinformatics 32: 1749–1751.

Noor, M. a. F., D. a. Garfield, S. W. Schaeffer and C. a. Machado, 2007 Divergence between the *Drosophila pseudoobscura* and *D. persimilis* genome sequences in relation to chromosomal inversions. Genetics 177: 1417–1428.

Obbard, D. J., J. J. Welch, K. W. Kim and F. M. Jiggins, 2009 Quantifying adaptive evolution in the *Drosophila* immune system. PLoS Genetics 5: e1000698.

Orr, H. A., 1995 Population genetics of speciation: the evolution of hybrid incompatibilities. Genetics 139: 1805–1813.

Orr, H. A., 2004 Dobzhansky, Bateson, and the genetics of speciation. Genetics 168: 1097–1104.

Orr, H. A., 2013 Haldane’s Rule. 28: 195–218.

Orr, H. A., L. H. Orr and H. A. Orri, 2013 Waiting for speciation : The effect of population subdivision on the time to speciation. Evolution 50: 1742–1749.

Palmieri, N., C. Kosiol and C. Schlötterer, 2014 The life cycle of *Drosophila* orphan genes. eLife 3: 1–21.

Patterson, J. T., 1947 The insemination reaction and its bearing on the problem of speciation in the mulleri subgroup. Univ Texas Publ 4720: 41–77.

Patterson, J. T., 1954 Studies in the genetics of Drosophila, pp.

Payseur, B. A., and L. H. Rieseberg, 2016 A genomic perspective on hybridization and speciation. Mol Ecol 25: 2337–2360.

Petrov, D. a., J. L. Schutzman, D. L. Hartl and E. R. Lozovskaya, 1995 Diverse transposable elements are mobilized in hybrid dysgenesis in Drosophila virilis. Proceedings of the National Academy of Sciences of the United States of America 92: 8050–8054.

Presgraves, D. C., 2007 Speciation genetics: Epistasis, Conflict and the Origin of Species. Current biology : CB 17: R125–127.

Presgraves, D. C., 2010 Speciation genetics: search for the missing snowball. Current biology : CB 20: R1073–1074.

R-Core-Team, 2013 R: A Language and Environment for Statistical Computing, pp. R Foundation for Statistical Computing, Vienna, Austria.

Rausch, T., T. Zichner, A. Schlattl, A. M. Stutz, V. Benes et al., 2012 DELLY: structural variant discovery by integrated paired-end and split-read analysis. Bioinformatics 28: i333–i339.

Ruan, J., and H. Li, 2020 Fast and accurate long-read assembly with wtdbg2. Nat Methods 17: 155–158.

Sackton, T. B., B. P. Lazzaro, T. A. Schlenke, J. D. Evans, D. Hultmark et al., 2007 Dynamic evolution of the innate immune system in *Drosophila*. Nature Genetics 39: 1461–1468.

Satyaki, P. R., T. N. Cuykendall, K. H. Wei, N. J. Brideau, H. Kwak et al., 2014 The Hmr and Lhr hybrid incompatibility genes suppress a broad range of heterochromatic repeats. PLoS Genet 10: e1004240.

Shultz, A., and T. B. Sackton, 2019 Immune genes are hotspots of shared positive selection across birds and mammals. eLife 8: e41815.

Simão, F. A., R. M. Waterhouse, P. Ioannidis, E. V. Kriventseva and E. M. Zdobnov, 2015 BUSCO: Assessing genome assembly and annotation completeness with single-copy orthologs. Bioinformatics 31: 3210–3212.

Smit, A. F. A., and R. Hubley, 2008 RepeatModeler Open-1.0.

Smit, A. F. A., and R. Hubley, 2013-2015 RepeatMasker Open-4.0, pp. RepeatMasker.

Stalker, H. D., and D. Streisinger, 1953 Taxonomy and Hybridization in the Cardini Group of Drosophila. 46: 343–358.

Thomas, P. D., M. J. Campbell, A. Kejariwal, H. Mi, B. Karlak et al., 2003 PANTHER: a library of protein families and subfamilies indexed by function. Genome Res 13: 2129–2141.

Turissini, D. A., J. A. McGirr, S. S. Patel, J. R. David and D. R. Matute, 2017 The Rate of Evolution of Postmating-Prezygotic Reproductive Isolation in Drosophila. Molecular Biology and Evolution 35: 312–334.

Unckless, R. L., V. M. Howick and B. P. Lazzaro, 2016 Convergent Balancing Selection on an Antimicrobial Peptide in *Drosophila*. Current Biology 26: 257–262.

Walker, B. J., T. Abeel, T. Shea, M. Priest, A. Abouelliel et al., 2014 Pilon: An integrated tool for comprehensive microbial variant detection and genome assembly improvement. PLoS ONE 9.

Welch, J. J., 2004 Accumulating Dobzhansky-Muller incompatibilities: reconciling theory and data. Evolution; international journal of organic evolution 58: 1145–1156.

Wickham, H., 2009 ggplot2: Elegant Graphics for Data Analysis.

Wigby, S., and T. Chapman, 2005 Sex peptide causes mating costs in female Drosophila melanogaster. Curr Biol 15: 316–321.

Wilder, J. a., and H. Hollocher, 2003 Recent radiation of endemic Caribbean *Drosophila* of the *dunni* subgroup inferred from multilocus DNA sequence variation. Evolution 57: 2566–2579.

Yang, Z., 2007 PAML 4: Phylogenetic analysis by maximum likelihood. Molecular Biology and Evolution 24: 1586–1591.

Zhang, H., P. Meltzer and S. Davis, 2013 RCircos : an R package for Circos 2D track plots.

Zhou, Q., and D. Bachtrog, 2015 Ancestral Chromatin Configuration Constrains Chromatin Evolution on Differentiating Sex Chromosomes in *Drosophila*. PLoS Genetics 11: 1–21.

Zhou, Q., H.-m. Zhu, Q.-f. Huang, L. Zhao, G.-j. Zhang et al., 2012 Deciphering neo-sex and B chromosome evolution by the draft genome of *Drosophila albomicans*. BMC Genomics 13: 109.

Zhou, W., F. Rousset and S. O’Neill, 1998 Phylogeny and PCR-based classication of *Wolbachia* strains using wsp gene sequences. Proc. R. Soc. 265: 509–515.

